# Sequestration of the phagocyte metabolite itaconate by *P. aeruginosa* RpoN promotes successful pulmonary infection

**DOI:** 10.1101/2025.08.07.669126

**Authors:** Ayesha Beg, Zihua Liu, Ying-Tsun Chen, Absar Talat, Griffin Gowdy, Jake Miller, Lindsey Florek, Lars Dietrich, Chu Wang, Ian Lewis, Tania Fok Lung Wong, Sebastian Riquelme, Alice Prince

## Abstract

The phagocyte immunometabolite itaconate, normally toxic to bacteria, functions as a signal to stimulate the adaptation of the pulmonary pathogen *Pseudomonas aeruginos*a to the lung. Itaconate is actively transported into *P. aeruginosa* where it induces σ^54^ *rp*oN expression and co-valently binds cysteine residues on RpoN. RpoN not only functions as a sink to limit itaconate toxicity but *S*- itaconated RpoN promotes increased utilization of the Entner Doudoroff pathway, optimizing bacterial metabolism in the setting of inflammation. *S*-itaconation of RpoN directs a global metabolic response that fuels pulmonary infection.

## Manuscript

*ESKAP*E pathogens such as *Pseudomonas aeruginosa*, whether antibiotic resistant or not, are a major cause of health care associated pneumonia world wide^1^. They are typically aspirated into the lung from abiotic sites such as puddles, showers or sinks and must rapidly adapt or risk immune clearance. To establish a nidus of infection, successful opportunists evade a variety of host antibacterial effectors. They must optimize metabolic activity selecting from the abundant carbon sources available but without fueling excessive growth that would elicit an immune response toxic to both host and pathogen ^2^. We wanted to understand how these bacteria sense they are in a human airway and coordinate gene expression to enable proliferation in response to this specific environmental challenge.

Both classical and alternative sigma factors coordinate bacterial sensing and adaptation to diverse environments. The alternative σ^54^ factor RpoN is among the over 400 *P. aeruginosa* transcriptional regulators that control bacterial gene expression in response to local cues. RpoN is involved in both the positive and negative regulation of genes that influence bacterial persistence in the respiratory tract, including multiple quorum sensing systems, biofilm formation, motility and especially metabolism ^3–6^. As RpoN directs pathways involved in the uptake and assimilation of desirable carbon sources to generate energy, it seemed likely that this transcription factor plays a major role in bacterial adaptation to the environment imposed by the infected airway.

Itaconate, synthesized by immunoresponsive gene 1 (*Irg1*), is a major airway metabolite released by phagocytes in response to infection ^7^. It modifies both host and bacterial proteins through covalent interactions with specific cysteine residues often altering metabolic activities ^8–10^. In the host Itaconate shapes immunoregulatory processes via *S*-itaconation of proteins involved in pro-inflammatory signaling, such as the NLRP3 inflammasome, the Nrf2-sequestrator Keap1 and succinate dehydrogenase.

In bacteria Itaconate is toxic ^11,12^ and acts as an electrophile inducing substantial membrane stress as well as blocking TCA cycle function and inhibiting carbohydrate catabolism particularly glycolysis^13,14^. As one of the most abundant metabolites in the infected airway, we postulated that itaconate serves as a host-specific signal that stimulates *P. aeruginosa* to adapt to the lung environment. In the experiments detailed in this report, we demonstrate how the binding of itaconate at two highly conserved cysteine residues in RpoN sequesters this electrophile and modifies RpoN function to promote *P. aeruginosa* carbohydrate catabolism fueling bacterial adaptation to the lung.

### RpoN directs *P. aeruginosa* responses to itaconate in the lung

*P. aeruginosa* pulmonary infection elicits a brisk phagocytic response, dominated by their abundant release of itaconate^14^ . To directly test the impact of RpoN dependent transcription on *P. aeruginosa in vivo*, we compared the outcomes of pulmonary infection in mice exposed to the WT PAO1 strain or a Δ*rpoN* mutant, appreciating that over 400 genes are under either positive or negative RpoN regulation^15^. The Δ*rpoN* strains achieved a higher bacterial load than the WT PAO1 (**Fig. 1a,b**) and stimulated slightly greater levels of IL-1α, IL-1β, MIG and MCP-1, but not IL-6 or TNFα (**Extended data Fig. 1a**) consistent with the increased bacterial load. Mice infected with WT PAO1 exhibited decreased amounts of itaconate in the airway (**Fig. 1c**) yet had similar numbers of phagocytes, the cells that produce itaconate (**Fig. 1d,e**). The apparent ability of WT bacteria to limit the accumulation of itaconate in the airways, could be either a bacterial or host effect. Taken together, these results suggested to us that RpoN may interact with itaconate specifically in its response to the airway metabolome.

**Fig. 1.**
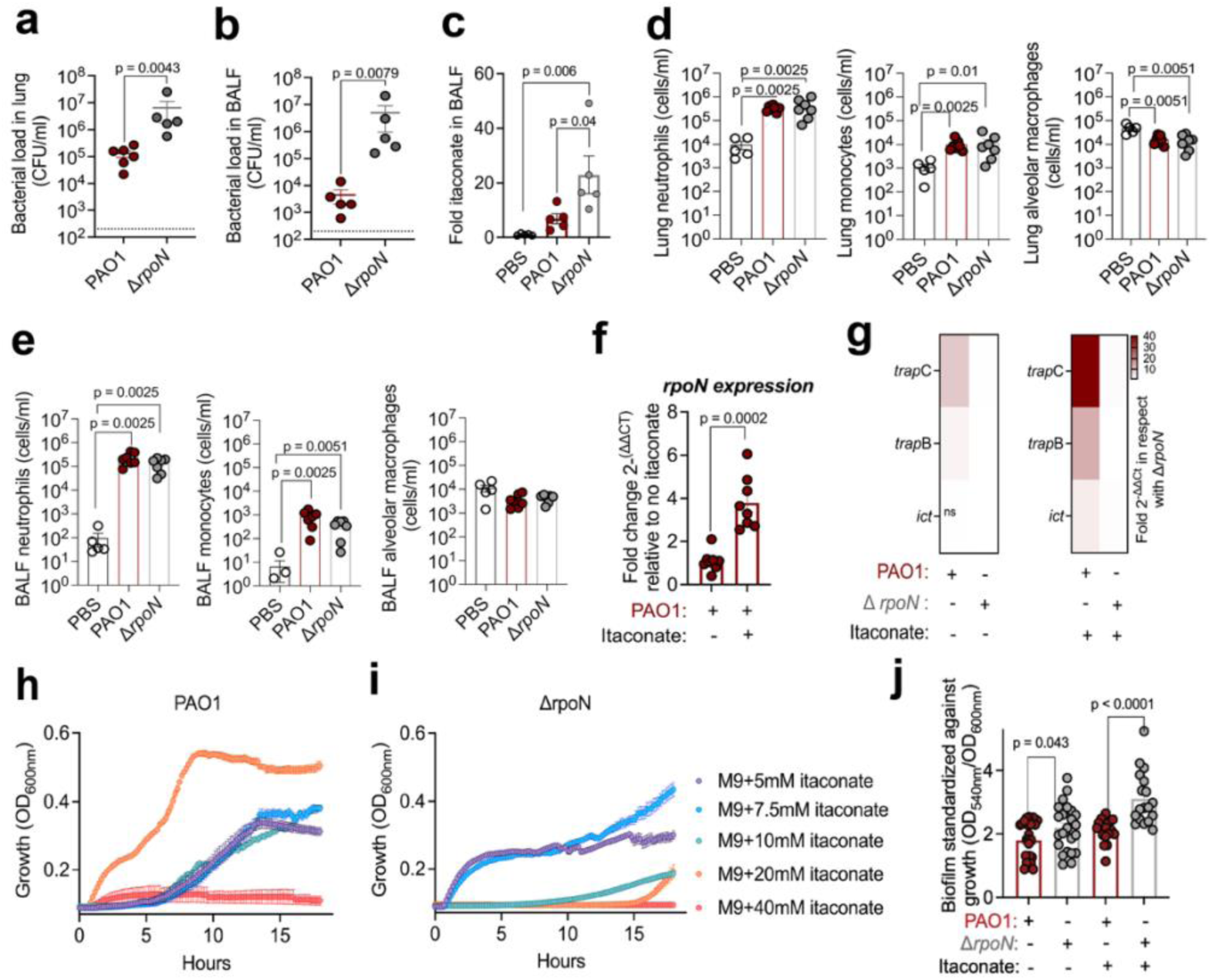
Expression of *rpoN* regulates bacterial response to pulmonary pathogenesis and itaconate. **a-e**, Characteristics of acute pulmonary infection in C57 BL/NJ6 mice following intranasal inoculation of WT or Δ*rpoN* PAO1 strains at 16 h post infection. Bacterial burden from (**a**) lung and (**b**) Broncheoalveaolar lavage fluid (BALF). (**c**) Relative abundance of itaconate in BALF. Immune cells in (**d**) BALF (**e**) lung. **f-j,** *In vitro rpoN-* mediated response to itaconate. RT-qPCR based quantification of (**f**) *rpoN* mRNA in PAO1 WT relative to itaconate and (**g**) itaconate metabolizing (*ict*) and transporter (*trap*B, *trap*C) genes in PAO1 WT relative to Δ*rpoN* +/- itaconate. Growth curves of (**h**) PAO1 and (**i**) Δ*rpoN* PAO1 strains in M9 minimal media supplemented with increasing concentrations of itaconate (5-40mM) as a sole carbon source. (**j**) Biofilm production (normalized to growth) by WT and Δ*rpoN* PAO1 strains in LB media +/- itaconate at 24 h. Data are presented as mean ± s.e.m.; *n* = 2 (**a,b**), *n = 2*(**c-g**), *n* = 2 (**h,i**) and *n* = 2 (**j**) replicates. Statistical significance was assessed unpaired Mann Whitney U *t*-test (**a,b,d-f**), Unpaired two-tailed Student’s *t*-test (**g**) and One way ANOVA using Tukey’s multiple comparison test with specific two-tailed unpaired *t*-test (**c,j**). **ns**(non-significant).

We observed that the organisms respond to itaconate by increasing *rpoN* mRNA expression, as well as expression of the itaconate transporters *trap*B, *trap*C and *ict* involved in itaconate assimilation^16^ (**Fig. 1f,g**); suggesting that the pathogen employs the σ^54^ factor as both sensor and defense mechanism in response to the immunometabolite. *In vitro* in the presence of itaconate *rpoN* expression protects *P. aeruginosa* proliferation (**Fig. 1h**). Whereas, the Δ*rpoN* mutant exhibits decreased growth rate (**Fig. 1i**) but increased production of biofilm in response to itaconate stress (**Fig. 1j**). Transcriptional studies confirmed that the Δ*rpoN* mutant responded to itaconate with increased expression of the families of genes involved in quorum sensing (*las*, *rhl)* and biofilm formation (*psl*,*pel*, alginate) mechanisms to protect against oxidant stress (**Extended data Fig. 1b**). These studies indicate the physiological relevance of RpoN for *P. aeruginosa* adaptation to an itaconate-rich environment like the lung. We next addressed how RpoN and itaconate might interact to regulate *P. aeruginos*a adaptation to the lung.

### Sequestration of itaconate via *S*-itaconation of RpoN enhances *P. aeruginosa* persistence in the lung

As an electrophile itaconate can function as a non-specific oxidant, but it also has a major role in post translational modification of specific targets by covalently binding available cysteine residues — i.e. S-itaconation; modifications that can result in either loss or gain of function^17^. The conserved cysteine residues in *P. aeruginosa* PAO1 RpoN at 218 and 275 are sites of *S*-itaconation identified using a biorthogonal probe (C3A) validated for itaconate (**Fig. 2a**, **Extended data Fig. 2a**)^10^. We postulated that *S*- itaconation of RpoN at these two cysteine residues is a key factor responsible for *P. aeruginosa* to thrive in the host lung.

**Fig. 2.**
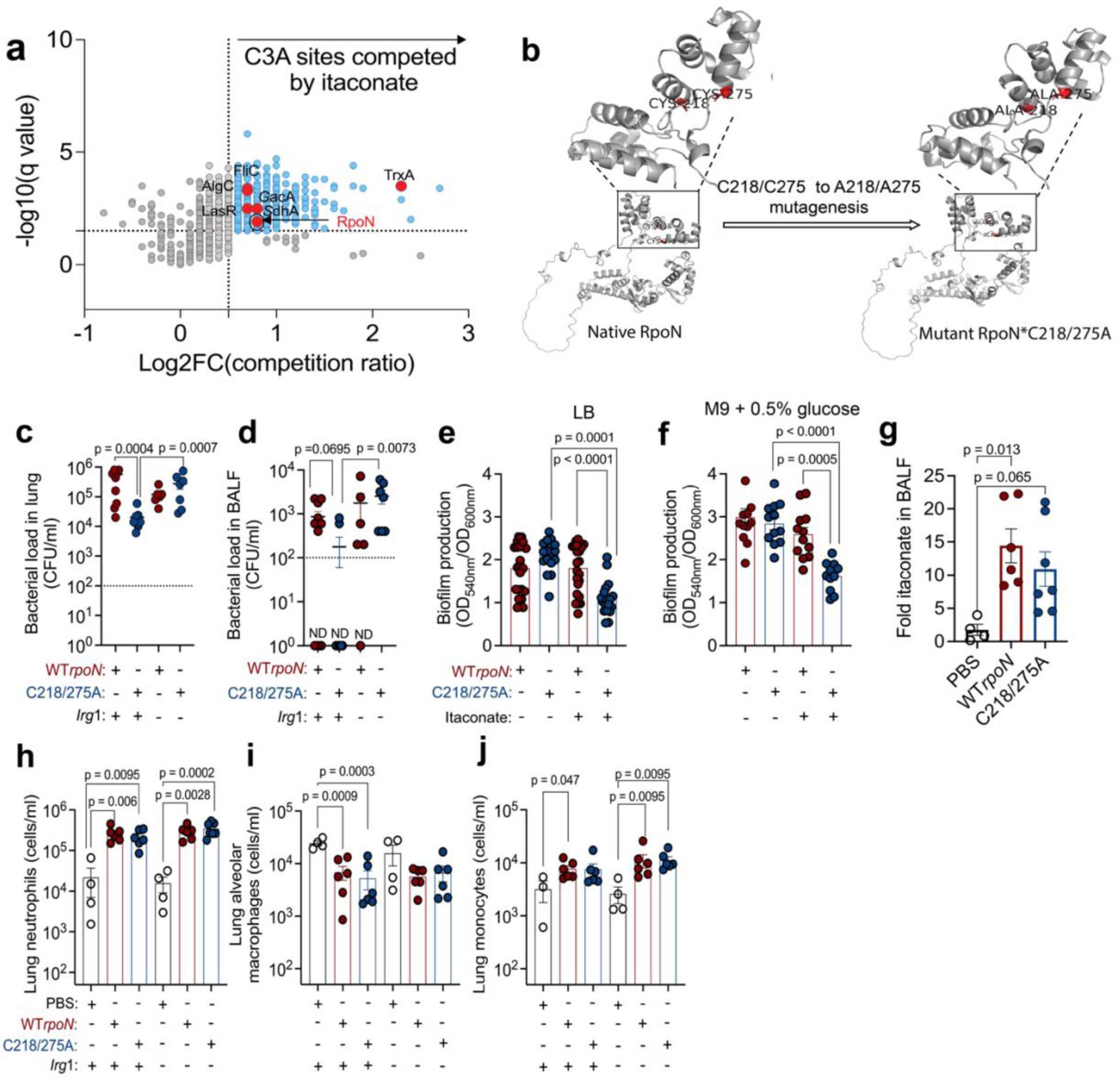
Impact of RpoN S-itaconation on *P. aeruginosa* pulmonary infection. (**a**) Identification of sites in live PAO1 WT that are labelled by C3A probe that are selectively competed by itaconate. Thresholds for the itaconate log2FC competition ratio and –log₁₀(q-value) are set at 0.5 and 1.5, respectively. Blue dots indicate sites increasingly competed by itaconate, while red dots highlight itaconated RpoN and common pathoadaptive proteins in *P. aeruginosa*. (**b**) Ribbon structure of RpoN highlighting substitution of C218 and C275 in native RpoN (red) to A218 and A275 in mutant RpoN (red). **c,d,g-j,** Characteristics of acute pulmonary infection in C57 BL/NJ6 WT or Irg1 ^-^/^-^ mice following intranasal inoculation of WT*rpoN* or C218/275A PAO1 strains at 16h post infection. Bacterial burden from (**c**) lung, (**d**) BALF. *In vitro* effects of itaconate on biofilm production (normalized to growth) by WT*rpoN* or C218/275A PAO1 in (**e**) LB media or (**f**) M9 media supplemented with 0.5% glucose at 24 h or 48 h, respectively. (**g**) Relative abundance of itaconate in BALF. (**h**) Neutrophils (**i**) alveolar macrophages (**j**) monocytes in lung. Data are presented as mean ± s.e.m.; *n* = 3 (**a**), *n =* 2 or 3(**c,d**), *n* = 2 (**e,f**) and *n* = 2 (**g-j**) replicates. Statistical significance was assessed using unpaired Mann Whitney U *t*-test (**c,d,h,i,j**) and One way ANOVA using Tukey’s multiple comparison test with specific two-tailed unpaired *t*-test (**e,f,g**).

We compared the outcomes of infection using *P.aeruginosa* strains with mutated *rpo*N lacking expression of the key cysteine residues that are itaconated versus the native *rpo*N. We complemented the PAO1 Δ*rpoN* mutant strain with a plasmid expressing either WT*rpoN* or an *rpoN* gene in which the cysteine residues at 218 and 275 were replaced with alanines and thus unable to be *S*-itaconated ^18^ (**Fig. 2b**, **Extended data Fig. 2b**). For simplicity of notation PAO1 Δ*rpoN* complemented with plasmid encoded WT *rpoN* is referred to as “WT*rpoN*” in contrast to PAO1 Δ*rpo*N expressing plasmid encoded C218/275A mutant *rpoN*, referred to as “C218/275A”. *P. aeruginosa* harboring RpoN available for *S*-itaconation, whether chromosomal or plasmid mediated had equivalent ability to sense the immunometabolite pressure *in vivo* and *in vitro*, as shown by similar levels of murine infection (**Extended data Fig. 2c**) and biofilm formation (**Extended data Fig. 2d**). We predicted that C218/275A lacking the cysteine residues essential for itaconate binding would exhibit biologically relevant phenotypes specifically in the presence of itaconate, but appreciated that other RpoN functions that are itaconate <u>independent</u> might be affected by the cysteine to alanine mutations. Phenotypic comparison revealed a slight growth advantage of the C218/275A mutant over WT*rpoN* when itaconate or succinate were used as sole carbon sources, whereas growth in glucose was inhibited by itaconate (**Extended data Fig.2 e-g**). Aside from these differences, both WT*rpoN* and C218/275A had equivalent rates of oxygen consumption and swarming motility, which are under RpoN regulation (**Extended data Fig. 2h-i**).

*In vivo*, RpoN *S*-itaconation supported *P. aeruginosa* survival in the respiratory tract. Compared with animals exposed to the C218/275A mutant, lungs of mice infected with the WT*rpoN* strain exhibited increased bacterial burden (**Fig. 2c,d**). This increased infection was dependent upon the presence of itaconate as both WT*rpoN* and the C218/275A mutant strains achieved similar levels of infection in *Irg*1-/- animals lacking itaconate (**Fig. 2c,d**). *In vitro*, the C218/275A mutant was significantly impaired in biofilm production in response to itaconate (**Fig. 2e,f**). Importantly, the bacteria expressing RpoN capable of sensing the itaconate via cysteine modification as well as those with an alanine substitution did not have significant alterations in other host effector responses against *P. aeruginosa*, such as their induction of itaconate production (**Fig. 2g**), stimulation of infiltrating phagocytes(**Fig. 2h,i,j**), or release of inflammatory cytokines (**Extended data Fig. 2j**). Together, these findings strongly suggest that itaconate sequestration by RpoN via *S*-itaconation confers *P. aeruginosa* with major survival advantages in a phagocyte dominated setting.

### RpoN *S*-itaconation promotes *P. aeruginosa* glucose catabolism and utilization of the Entner Doudoroff Pathway

We anticipated that the protection conferred to *P. aeruginosa* by RpoN *S*itaconation in the lung would involve mechanisms that promote establishment of the bacterial biomass in the lung. Transcriptomic profiles of the WT*rpoN* and C218/275A mutant (**Supplementary Table 1**) indicated several changes attributable to the C-A mutations under control conditions (**Extended data Fig. 3a,c**), as well as marked differences in key pathways in the presence of itaconate (**Fig. 3a, Extended data Fig. 3b**). While itaconate itself can be assimilated by *P.aeruginosa*^14^, its preferred mechanism to generate ATP and other precursor molecules under conditions of oxidant stress is via the energetically efficient Entner Doudoroff (ED) pathway (**Fig. 3a**) ^19–22^. This prioritizes the allocation of glucose to a range of metabolic networks involved in the synthesis of antioxidants, such as NADP+, as well as anthranilate metabolism, quorum sensing, iron scavenging and nucleotide synthesis all important for biomass expansion. We observed that itaconated RpoN was associated with significantly increased expression of *gcd* which converts glucose to gluconate as compared with the C218/275A mutant, upregulation of genes associated glucose oxidation to ketogluconates (*kgu*DET) and their utilization through the ED Pathway (*zwf*, *pgl*, and *eda)* (**Fig. 3c-e**). There was also increased expression of the antranilates, *ant*ABC, which participate in the degradation of tryptophan via the kyrenurine pathway which generates NAD^+^ and NADP^23,24^, cofactors needed to accept electrons in the production of gluconate and glycerol-3 phosphate via DHAP (**Fig. 3a,b**).

**Fig. 3.**
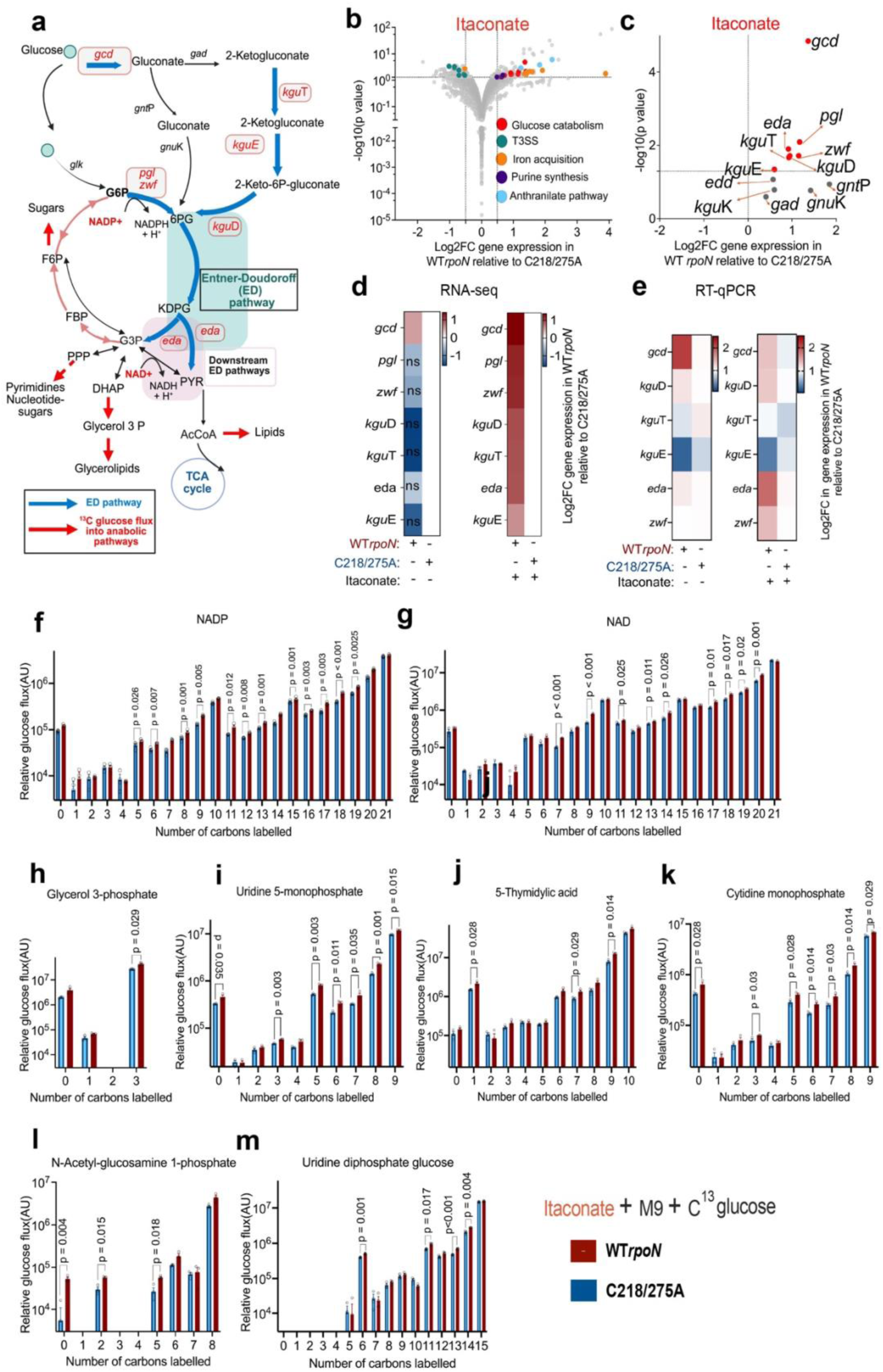
Impact of RpoN S-itaconation on transcriptional profile and anabolic glucose utilization in *P. aeruginosa*. (**a**) Schematic of glucose metabolism in *P. aeruginosa*, highlighting increased expression of genes (red) involved in the gluconate and Entner-Doudoroff (ED) pathways (blue), as well as enhanced glucose flux into anabolic product formation (red arrows) from primary metabolic pathways in WT *rpoN* versus C218/275A strains grown with itaconate. (**b**) Bulk RNA seq to determine differentially expressed genes in WT *rpoN* relative to C218/275A grown in LB media with itaconate. Grey dots represent all genes; colored dots represent genes of specific pathways. Y axis: logarithmic scale, cut off set on Y axis: -log10 (*p* value) ≥ 1.3; cut off set on X axis Log2FC(Fold change) > 1. (**c**) Volcano plot of differentially expressed genes specific to gluconate or ED pathway in presence of itaconate. Significantly expressed genes associated with gluconate and ED pathways as determined by (**d**) RNA seq and (**e**) RTqPCR in WT*rpoN* relative to C218/C275A with or without itaconate. Effect of itaconate on the ^13^C glucose carbon abundance in different isotopologues of metabolites in WT*rpoN* relative to C218/C275A. Enrichment of ^13^C glucose into isotopologues of (**f, g**) cofactors: NAD and NADP, (**h**) glycerol, (**i- k**) pyrimidines: UMP, CMP, dTMP, and (**l, m**) sugars. Data are presented as mean ± s.e.m.; *n* = 2 (**b,c**), *n* = 2 (**d,e**), *n =* 3 (**f-m**) replicates. Significance is determined by Wald *t* test(**b,c,d**) and Unpaired two-tailed Student’s *t*-test(**e,f-m**). **ns**(non-significant). Schematics in (**a**) is created by BioRender (https://biorender.com).

Using carbon tracing assays, we corroborated how RpoN *S*-itaconation facilitated *P. aeruginosa* [^13^C]-glucose catabolism comparing the effects of itaconate on WT*rpoN* and C218/275A mutants (**Supplementary Table 2**). In the presence of itaconate, more of the [^13^C] signature was integrated into specific isotopologues of NADP+, NAD+ and glycerol -3- phosphate (**Fig. 3f,g,h**) core elements on the ED pathway. There was significant [^13^C] enrichment in nucleic acid precursors; uridine monopohosphate (UMP), 5-thymidylic acid (dTMP) and cytidine monophosphate (CMP) in WT*rpoN* as compared to C218/C275A as well (**Fig. 3i-k**). WT*rpoN* incorporated significantly more [^13^C] into lipid and rhamnolipid precursors derived from acetylCoA, such as 2-hydroxycaproic acid and D-2Hydroxyglutaric acid (**Extended data Fig. 3g**) and into the glycerolipid precursors glycerol-3 phosphate and O-phosphoethanolamine (**Fig. 3h, Extended data Fig. 3g**). WT*rpoN* also incorporated more [^13^C]-glucose into *N*-acetyl-glucosamine-1-phosphate (GlcNAc-1-P) and UDP-glucose (UDP-Glc)(**Fig. 3l,m**), essential components of peptidoglycan and biofilm.

We also observed changes in ^13^C glucose flux attributed solely to C-A mutations (**Extended data Fig. 3e,f)**; namely decreased ^13^C enrichment into guanine diphosphate, ornithine, alanine, asparagine and allantoin (**Extended data Fig. 3h**), but increased ^13^C incorporation into L-cystathione and niacinamide (**Extended data Fig. 3i)**. The ^13^C incorporation into GlcNAc-1-P and UMP was slightly reduced under basal conditions in WT*rpoN* organisms decreased significantly upon itaconate exposure (**Extended data Fig. 3j**). Overall the transcriptomic and carbon tracing studies demonstrate that the availability of itaconate to modify RpoN provides a coordinated enhancement of the fundamental metabolic activities required for bacterial proliferation, promoting successful *P. aeruginosa* infection in a setting of inflammation and oxidant stress.

### The conservation of RpoN cysteine residues is a feature of successful *P. aeruginosa* pulmonary infection

Our findings thus far indicate that post translational modification of RpoN by itaconate enhances the ability of the organisms to establish pulmonary infection by optimizing bacterial metabolic activity, as modeled by the laboratory standard strain PAO1. Although *rpoN* is an essential gene^15^, it is a site of mutation in clinical isolates ^25^. We postulated that the cysteine residues at 218 and 275 in RpoN would be highly conserved in *P. aeruginosa* from patients with pneumonia if they are biologically important to enable infection amidst phagocytes and itaconate stress. We analyzed *rpoN* sequences in publically available data sets from cinical isolates of *P. aeruginosa*, known to associated with either acute ICU or chronic (cystic fibrosis (CF) pulmonary infection (**Fig. 4a**). We identified several mutations in the clinical isolates, mostly synonymous variants, a few missense mutations or deletions. The frequency of mutations was significantly greater in isolates from long standing CF infection than in the ICU isolates (**Fig. 4b**). A comparison of the *rpoN* genomi*c* sequences and predicted proteins from these different datasets indicated conservation of the cysteine residues in the same location as in the reference PAO1 sequence (**Fig. 4c**). This is a region of RpoN expected to be involved in RNA polymerase binding^18^, outside of the canonical DNA binding domain associated with the σ^54^ -12 and -24 regions of target genes (**Fig. 4d,e**). As depicted in the ribbon diagrams, mutations in the clinical strains did not appear likely to involve critical regions of RpoN, those involved in RNAP interactions or DNA recognition sites **(Fig. 4d**), although they could affect interactions with enhancer binding proteins^26^. The conservation of cysteines at positions 218 and 275 across the *P. aeruginosa* isolates from pneumonia patients worldwide is consistent with the physiologic role of S-itaconation of RpoN to promote the pathogenesis of *P*. *aeruginosa* pulmonary infection.

**Fig. 4.**
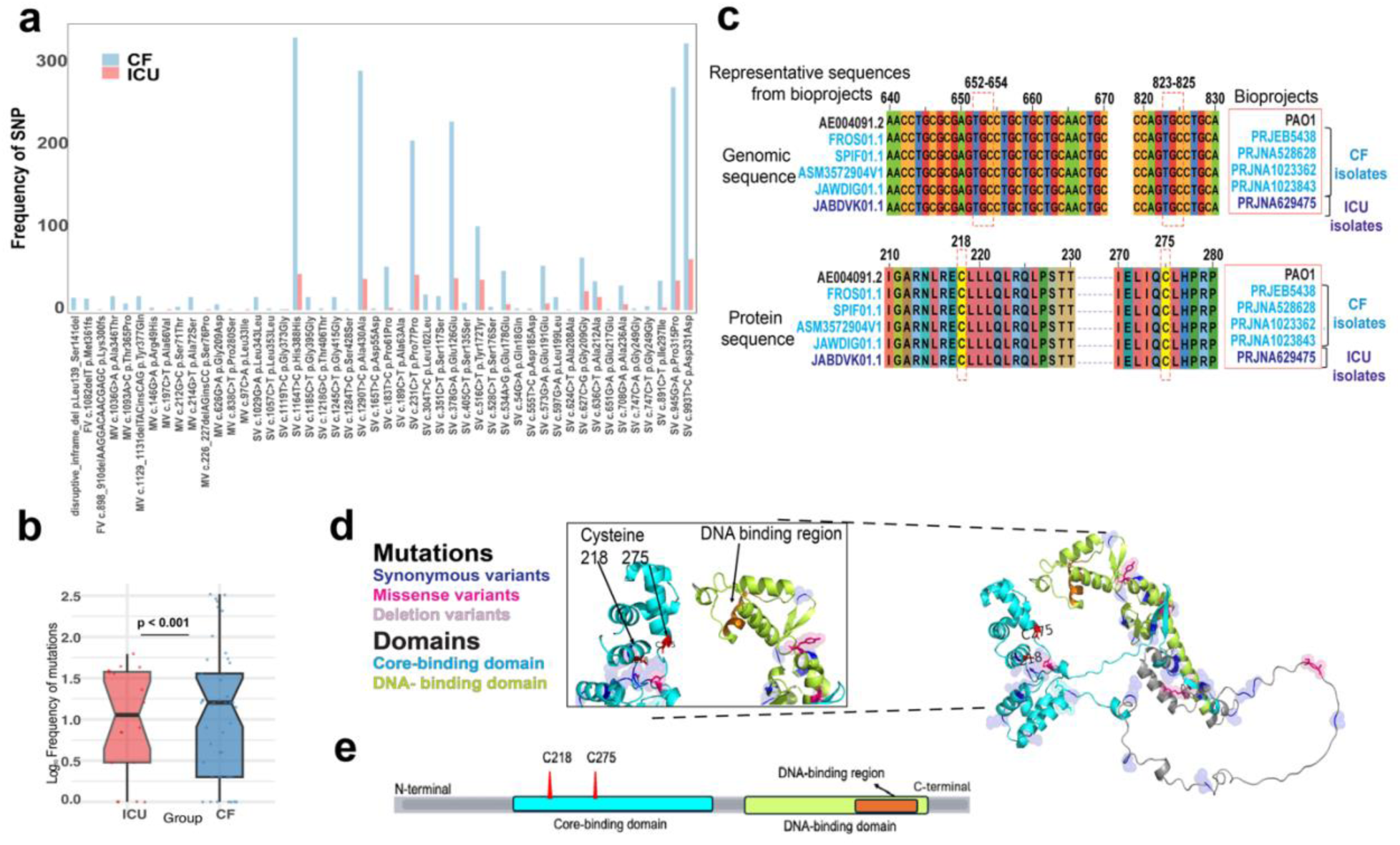
Clinical isolates conserve RpoN cysteines modified by itaconate (**a**) Relative frequency of mutations in the *rpoN* gene of *P. aeruginosa* strains from CF (n = 374) and ICU patients (n = 62) isolates (SV: synonymous variant; MV: missense variant). (**b**) Box-plot representing significant difference in frequency of total mutations in *rpoN* gene among CF and ICU groups. (**c**) Alignment of *rpoN* sequences from representative datasets (labeled by source BioProject number) highlighting the conservation of cysteines at positions 218 and 275, relative to PAO1, at both genomic (652–654; 823–825) and protein (218/275) sequence levels. (Bioprojects: CF-blue; ICU: purple). (**d**) Ribbon structure of RpoN protein illustrating different mutations and domains, **(inset)** position of conserved cysteines 218/275 relative to DNA binding region of RpoN. (**e**) Linear map of RpoN protein from N-C terminus, depicting positions of conserved cysteines, DNA binding region, core- and DNA- binding domain. Significance is determined by Shapiro-Wilk normality test(**b**).

## Discussion

The goal of this study was to establish how *P. aeruginosa* senses and responds to aspiration into the lung, an oxidant rich environment replete with phagocytes and their products. Aided by decades of accumulated data characterizing the metabolic and genetic properties of this ubiquitous pathogen, we identified the alternative σ^54^ factor RpoN as highly responsive to the phagocyte-specific immunometabolite itaconate, which seems especially biologically appropriate. Whereas the importance of itaconate in regulating innate immune responses to pathogens is increasingly appreciated ^7,8,14^, its impact on bacteria as one of the most abundant metbolites in the infected airway, has been less well appreciated. Upon sensing itaconate, genes directing its transport and assimilation, as well as *rpo*N expression itself, are all upregulated. We found that *P. aeruginosa* detoxifies itaconate by covalent binding to cysteine residues of RpoN and that itaconated RpoN boosts bacterial metabolism by promoting use of the preferred Entner Doudoroff Pathway to generate ATP. The itaconated transcription factor also enhances the production of NAD and NADP co-factors utilized in this pathway by increasing expression of the anthranilate-tryptophan-kyrenurine cascade. Thus WT *P. aeruginosa* responds to the challenges of itaconate through effects on a global transcription factor. It is striking, but not surprising, that a host-specific immunometabolite has such a major impact promoting *P. aeruginosa* infection.

Our data emphasize the importance of *rpoN*, a bacterial gene not typically considered a virulence factor in the pathogenesis of pulmonary infection. The expression of *rpoN* contributes to the exceptional metabolic versatility of *P. aeruginosa*. While the pathogens can metabolize itaconate, the S- itaconation of RpoN confers more pervasive benefits to support persistence in the oxidant-rich lung.

Itaconated RpoN by promoting the generation and utilization of gluconate, which accumulates in cystic fibrosis patients with pulmonary infection^20^, optimizes ATP production in settings in which glucose is limited such as the airway^27^, a likely consequence of substrate competition from neutrophils ^22,28^.

Utilization of the Entner-Doudoroff pathway and production of NADP also increases protection from oxidants via glutathione and generates less ROS itself than glycolysis. Thus, as an in vivo response to the challenges of survival in the lung, the ability of these pathogens to exploit host itaconate helps to explain their success as opportunistic pathogens. As other Gram-negative bacteria and members of the *ESKAPE* family ^29^ express *rpoN* with conserved cysteine residues in a similar configuration as *P. aeruginosa*, strategies to prevent bacterial adaptation to the itaconate dominated milieu of the infected lung by interfering with RpoN might prove more useful than the development of vaccines targeting individual gene products^30^.

## METHODS

### Key Reagents

Dream Taq PCR mastermix (K1081, Thermo Scientific™), EcoRI-HF(NEB R0101S) and HindIII-HF(NEB, R3104S), QIAprep Spin Miniprep Kit (QIAGEN,27104), T4 DNA Ligase (5 U/μL)(Thermo Scientific™, EL0011), ReliaPrep™ DNA Clean-Up and Concentration System (Promega A2893).

Tetracycline hydrochloride (Sigma, T7660), Itaconate (Sigma# I29204), D-(+)-Glucose (Sigma-Aldrich, G8270) M9 minimal media (Gibco™A1374401), Seahorse calibrant (Agilent #100840-000).

RNAprotect Bacteria Reagent (QIAGEN, 76506), Proteinase K (QIAGEN, 19131), TRK lysis buffer (Omega Bio-tek, R6834-02), Multiscribe Reverse Transcriptase (Applied Biosystems, 43-688-14), PowerUp SYBR Green Master Mix (Applied Biosystems, 25742), DNA-free™ DNA Removal Kit (Invitrogen, AM1906).

Antibodies and staining reagents: Anti-CD45-AF700 (BioLegend, 103127), anti-CD11b-AF594 (BioLegend, 101254), anti-CD11c-BV605 (BioLegend, 117334), anti-SiglecF-AF647 (BD, 562680), anti-MHCII-APC-Cy7

(BioLegend, 107628), anti-Ly6C-BV421 (BioLegend, 128032), and anti-Ly6G-PerCP-Cy5.5 (BioLegend, 127616), LIVE/DEAD viability dye (Invitrogen, L23105A), 15.45 μm DragonGreen, Bangs Laboratories Inc., FS07F).

### Mouse experiments

All animal experiments were conducted in accordance with institutional guidelines at Columbia University Irving Medical Center and approved under IACUC protocol AABD5602. Wild-type (WT) C57BL/6NJ mice (7–8 weeks old, 20–25 g; Jax #005304) were obtained from the Jackson Laboratory.

Irg1⁻/⁻ (Acod1⁻/⁻) mice (Jax #029340) were also obtained from the Jackson Laboratory and bred in-house at Columbia University Irving Medical Center. Both WT and Irg1⁻/⁻ mice were immunocompetent and did not receive any medical or drug treatments prior to infection. Each in vivo experiment included an equal number of male and female mice (50:50 ratio), and no sex-based differences were anticipated. Animals were randomly assigned to cages and housed in barrier facilities under standard conditions (12-hour light/dark cycle, temperature 18–23 °C, 30–50% humidity). Mice were fed a regular irradiated chow diet (Purina Cat #5053, distributed by Fisher).

### Growth condition of bacterial strains

*Pseudomonas aeruginosa* strains – PAO1, Δ*rpoN*, Δ*rpoN* :p*rpoN*, Δ*rpoN* :pC218/C275, Δ*rpoN* empty vector control (EVC) were used pHERD26T . In the experiments, laboratory *Pseudomonas* strains were grown in LB for overnight and subcultured until exponential phase. All strains were streaked and maintained on LB agar plates, while Δ*rpoN* :p*rpoN*, Δ*rpoN* :pC218/C275, Δ*rpoN*(EVC) were maintained on LB agar plates supplemented with 50 µg/mL of tetracycline.

### Strains and plasmids

Strains, plasmid and primers used in this study are listed in **Supplementary Table 3.** We constructed an *rpoN* mutant in which the codons for cysteines at positions 218 and 275 were substituted with alanine using the overlap extension PCR method^1^. The *rpoN* sequence was PCR-amplified using

Dream Taq PCR mastermix (Thermo Scientific™, K1081) from genomic DNA of PAO1WT using primer pairs, RpoN_F and RpoN_R. The mutation at 218 position was introduced using primer sets RpoN_F

/218_R and 218_F/ RpoN_R, followed by fusion of resultant PCR products using RpoN_F and RpoN_R. Similarly, substitution mutation was introduced in C218A\**rpoN* at 275 position using primer set

RpoN_F/275_R and 275_F/RpoN_R, followed by fusion of resultant PCR products using RpoN_F and RpoN_R. The amplicons, *rpoN* and C218/275A\**rpoN* were cloned into vector pHerd26T using EcoRI-HF(NEB R0101S) and HindIII-HF(NEB, R3104S), yielding plasmids pHerd26T-WT*rpoN* and pHerd26T- C218/C275A. Verified plasmids were electroporated into Δ*rpoN* PAO1, and Transformants were selected on LB agar plates supplemented with 50 µg/mL of tetracycline, as previously described^2^. Clones were screened by PCR using flanking primers RpoN_F and RpoN_R, followed by Sanger sequencing of the amplicons.

### Mouse pneumonia model

Mouse pneumonia infection was carried out with 7-8 weeks old male and female (C57BL/NJ6 WT and *Irg1^−/−^* (*Acod1^−/−^*) mice as previously described ^3^. After anesthesia, animals were exposed to either PBS, WT PAO1, D*rpoN*, D*rpoN* :p*rpoN*, or D*rpoN* :pC218/C275. A 50µL volume of ∼10^6^ <u>CFUs</u> of either *P. aeruginosa* strain or PBS alone (non-infected) was intranasally inoculated into the mice. Mice were euthanized at 16h post infection; whole lungs and BALF were collected aseptically. The lung was homogenized through 40 µm cell strainers (Falcon, 352340). Aliquots of BALF and lung homogenates were serially diluted and plated on LB agar plates (supplemented with 50 µg/mL tetracycline when needed) to determine the bacterial burden. The BALF and lung homogenates were spun down and the BALF supernatant was collected for cytokine and untargeted metabolomic analysis. After hypotonic lysis of the red blood cells, the remaining BALF and lung cells were prepared for fluorescence-activated cell sorting (FACS) analysis as described below. No calculation was used to determine the number of mice required. No data blinding was performed

### Flow cytometry for immune cell recruitment in infection studies

Immune cell profiling in BALF and lung tissue was performed by staining isolated cells with a LIVE/DEAD viability dye (Invitrogen, L23105A) and a fluorescent antibody cocktail, along with 10 µL of counting beads (15.45 μm DragonGreen, Bangs Laboratories Inc., FS07F). The antibody panel included: anti-CD45-AF700 (BioLegend, 103127), anti-CD11b-AF594 (BioLegend, 101254), anti-CD11c-BV605 (BioLegend, 117334), anti-SiglecF-AF647 (BD, 562680), anti-MHCII-APC-Cy7 (BioLegend, 107628), anti- Ly6C-BV421 (BioLegend, 128032), and anti-Ly6G-PerCP-Cy5.5 (BioLegend, 127616), each diluted 1:200 in PBS. Staining was carried out for 1 hour at 4 °C. After washing, cells were fixed in 2% paraformaldehyde (Electron Microscopy Sciences, 15714-S) and analyzed on a BD LSRII flow cytometer using FACSDiva v9 software. Data were processed with FlowJo v10.Cell populations were identified based on the following markers: **Alveolar macrophages:** CD45⁺ CD11b⁺/⁻ SiglecF⁺ CD11c⁺; **Neutrophils:** CD45⁺ CD11b⁺ SiglecF⁻ MHCII⁻ CD11c⁻ Ly6G⁺ Ly6C⁺/⁻; **Monocytes:** CD45⁺ CD11b⁺ SiglecF⁻ MHCII⁻ CD11c⁻ Ly6G⁻ Ly6C⁺/⁻.

### Cytokine analysis

Cytokine levels in mouse BALF supernatants were measured by Eve Technologies (Calgary Canada) using bead-based multiplex technology.

### Untargeted metabolomic analysis

Metabolite profiling of BALF samples was conducted using high-resolution LC-MS at the Calgary Metabolomics Research Facility (Calgary, Canada). Metabolite extraction was performed using a 50% methanol (Supelco #106018) and water (v/v) solution. LC-MS analysis was carried out on a Q Exactive HF Hybrid Quadrupole-Orbitrap mass spectrometer (Thermo Fisher Scientific) coupled to a Vanquish UHPLC system (Thermo Fisher Scientific). Chromatographic separation was conducted on a Syncronis HILIC UHPLC column (2.1 mm × 100 mm × 1.7 μm, Thermo Fisher Scientific) using a binary solvent system at a flow rate of 600 µL/min. Solvent A was 20 mM ammonium formate at pH 3 in mass spectrometry-grade water, and solvent B was mass spectrometry-grade acetonitrile containing 0.1% formic acid (v/v). The gradient program was as follows: 0–2 min, 100% B; 2–7 min, 100%–80% B; 7–10 min, 80%–5% B; 10–12 min, 5% B; 12–13 min, 5%–100% B; 13–15 min, 100% B. A 2 ml of sample was injected. The mass spectrometer operated in negative full-scan mode at a resolution of 240,000, scanning from 50–750 m/z. Metabolites were identified by matching observed m/z values (±10 ppm) and retention times with those of commercial metabolite standards (Sigma-Aldrich). Data analysis was conducted using E-Maven v0.10.0.

### Isolation of bacterial RNA

WT*rpoN* and C218/C275 strains were grown overnight in LB media (+50 µg/mL tetracycline). Overnight cultures were inoculated (1/100) into LB (+50 µg/mL tetracycline) with or without itaconate (Sigma# I29204) and were grown at 37 °C to late exponential phase. Approximately, 2 × 10⁸ *P. aeruginosa* cells were harvested by centrifugation, and the resulting pellet was resuspended in RNAprotect Bacteria Reagent (QIAGEN, 76506). The suspension was vortexed briefly and incubated at room temperature (RT) for 10 minutes before centrifugation. The bacterial pellets were then lysed in a buffer pH 8.0 (containing 30 mM Tris (Corning, 46-030-CM), 1 mM EDTA (Thermo Fisher Scientific, 1861283), 15 mg/mL lysozyme (Sigma, L6876), and 200 mg/mL proteinase K (QIAGEN, 19131)), with incubation at RT for 10 minutes. Following lysis, TRK lysis buffer (Omega Bio-tek, R6834-02) and 70% ethanol (v/v) were added to the mixture. The lysates were then applied to E.Z.N.A. RNA isolation columns (Omega Bio-tek), and total RNA was extracted according to the manufacturer’s protocol. Residual genomic DNA was removed by DNase treatment using the DNA-free™ DNA Removal Kit (Invitrogen, AM1906).

### Complementary DNA (cDNA) synthesis and qRT–PCR

Complementary DNA (cDNA) was synthesized from RNA using Multiscribe Reverse Transcriptase (Applied Biosystems, 43-688-14). Quantitative real-time PCR (qRT-PCR) was carried out with gene-specific or housekeeping primers as listed in **Supplementary Table 3**, PowerUp SYBR Green Master Mix (Applied Biosystems, 25742), and the StepOnePlus Real-Time PCR System (Applied Biosystems). Relative gene expression levels were calculated using the ΔΔCt (delta-delta Ct) method.

### Bulk RNA sequencing

Bacterial RNA was extracted as previously described. A ribosomal RNA (rRNA)-depleted cDNA library was prepared according to the manufacturer’s instructions using the Universal Prokaryotic RNA-Seq Prokaryotic AnyDeplete kit (NuGEN #0363-32) and sequenced with Illumina HiSeq. Raw base call files were converted to fastq format using Bcl2fastq. Quality-filtered reads were then aligned to the *P. aeruginosa* PAO1 reference genome (RefSeq: GCF_000006765.1) using STAR Aligner v2.7.3a. Aligned reads were annotated with read group information and duplicate reads were identified using Picard Tools v2.22.3. Quantification of raw counts was performed using FeatureCount**s** from the Subread package v1.6.3. Differential gene expression analysis was conducted using DESeq2 in R v3.5.3 and low abundant transcript having threshold of 3 reads per gene were excluded in the analysis. The bacterial transcriptional data were plotted as a volcano plot heatmap plot using GraphPad Prism (v10.0c).

### 13C glucose labeling and stable isotope tracing

WT*rpoN* and C218/C275 strains were grown overnight in LB media (+50 µg/mL tetracycline) with or without 20mM of itaconate. The overnight cultures were washed and resuspended in the same volume of M9 minimal media (Gibco™A1374401). The cultures were inoculated (1/50) into M9 minimal media supplemented with 7.5 mM of ^13^C glucose (Sigma #389374) (+50 µg/mL tetracycline) and grown at 37 °C to late exponential phase. For metabolite extraction, each culture was pelleted and washed with PBS twice centrifuged in 2000 g for 10 minutes at 1 °C. The pellets were resuspended in a 3:1 methanol:water extraction solution and lysed with 10 freeze-thaw cycles by alternating emersion in liquid nitrogen and a dry-ice/ethanol bath. The debris was removed by centrifugation at 14,000 xg for 5 min at 1 °C and the supernatant was stored for analysis. Targeted LC/MS analysis was performed on a Q Exactive Orbitrap mass spectrometer (Thermo Scientific) coupled to a Vanquish UPLC system (Thermo Scientific). The Q Exactive operated in polarity-switching mode. A Sequant ZIC-HILIC column (2.1 mm i.d. × 150 mm, Merck) was used for separation of metabolites. Flow rate was set at 150 μL/min. Buffers consisted of 100% acetonitrile for mobile A, and 0.1% NH_4_OH/20 mM CH_3_COONH_4_ in water for mobile B. Gradient ran from 85 to 30% A in 20 min followed by a wash with 30% A and re-equilibration at 85% A.

Metabolites were identified based on exact mass within 5 ppm and standard retention times. Relative quantitation was performed based on peak area for each isotopologue. All data analysis was done using MAVEN 2011.6.17.

### *In situ* itaconation profiling of cysteines in *P. aeruginosa*

Cysteines in proteome subjected to S-itaconation were identified using the workflow described here^4^. *P. aeruginosa* PAO1 were grown in LB media at 37 °C to stationary phase. Cultures were washed and resuspended in pre-chilled PBS. Where 800 μL of *P. aeruginosa* PAO1 suspension in PBS was treated without or with 80 μL pH-adjusted itaconate was incubates on the ThermoMixer (950 rpm, 1 h, 37°C, 30 min). Both groups were then incubated with 8 μL of the C3A probe (950 rpm, 37 °C, 1 h). Cells were pelleted, washed, resuspended in 1 mL of 0.1% PBST, and lysed. Lysates were clarified by centrifugation, and the supernatant was transferred to fresh tubes. After competition, both groups were incubated with 8 mL of the C3A probe on the ThermoMixer (950 rpm, 37 °C, 1h). Bacteria were pelleted, washed and resuspended in 1 mL of 0.1% PBST for lysis. Lysates was centrifuged (20,000 g, 10 min, RT) to remove the debris and the supernatants was transferred into a new tubes.Click reaction was carried out at 29 °C for 1 h (1200 rpm)on a Thermomixer with 106 μL of Click reagent mix comprising of : 60 μL TBTA ligand, 20 μL 50 mM CuSO₄, 20 μL freshly prepared 50 mM TCEP, and 6 μL 20 mM acid-cleavable azide-biotin tag (Confluore). The click-labeled lysates were precipitated by 5 mL of MeOH/chloroform (4:1) and 3 mL of ddH2O. Precipitates were washed twice with pre-chilled MeOH, resuspended in 1 mL of 1.2% SDS in PBS, followed by sonication, heating for 10 min at 90 °C and centrifuged (20,000 g, 10 min, RT) to remove excessive copper. Supernatants were combined with the pre-washed streptavidin beads (Thermo Fisher) and incubated for 4 h at 29°C. Beads were washed PBS three times and ddH2O three times, followed by resuspension in 500 mL of 8 M urea (Sigma)/100 mM TEAB (Sigma). The samples were reduced on ThermoMixer (1,200 rpm, 30 min, 37°C) by 25 mL of 200 mM DTT (Shanghai Yuanye Bio-Technology Co., Ltd) and alkylated on the ThermoMixer (1,200 rpm, 30 min, 35°C) by 25 mL of 400 mM 2-iodoacetamide (Sigma). The beads were then resuspended in resuspended in 200 mL of 2 M urea/100 mM TEAB containing 1 mM CaCl2 and 10 ng mL1 trypsin (Promega) for digestion on ThermoMixer (1,200 rpm, 30 min, 35C). Post-digestion the supernatants were carefully transferred into a new 1.5 mL Protein LoBind Tubes (Eppendorf). The beads were washed, and post-washed supernatants were pooled with primary digest. The digested peptides were subjected to mass spectrometry for identification of itaconation of cysteines.

### Genomic analysis of clinical isolates

Publicly available whole genome sequences(WGS) for *Pseudomonas* CF isolates (Bioproject accession number : PRJEB5438, PRJNA528628, PRJNA1023362,PRJNA1023843 on NCBI) and ICU isolates (ENA accession number : PRJNA629475) were used for mutation analysis with respect to the reference genome PAO1(Genomics accession no.: AE004091.2).SNPs in *rpoN* were identified in CF and ICU isolates of *Pseudomonas aeruginosa* using Snippy v4.6.0 (https://github.com/tseemann/snippy), with the PAO1 genome as the reference. The detected variants, including SNPs and indels, were visualized using Jalview for comparative sequence analysis to determine the conservation of cysteines residues at the codon and protein sequence level. Representative sequences of *rpoN* from each bioprojects with respect to reference PAO1 were used to represent conservation of cysteine residues.

### Extracellular flux analysis

WT*rpoN*, C218/C275 and Δ*rpoN* strains were grown in LB overnight, then washed twice with filtered PBS before inoculating (1 × 10^7^) into 450 μl of 1X of M9 minimal media supplemented with 2 mM of MgSO_4_ and 0.1 mM of CaCl_2_ in Seahorse XFe24 cell culture plate (Agilent #102334-000B). 50 μl of 75 mM glucose or 200mM itaconate in M9 media was loaded onto the Seahorse XFe24 cartridge, which was previously hydrated in Seahorse calibrant (Agilent #100840-000) overnight at 37 °C. After 60 minutes of stabilization at the basal state, glucose or itaconate was acutely injected into the plate to achieve the final concentration of 7.5 mM or 20mM, respectively. Bacterial extracellular acidification and oxygen consumption rates were captured with Seahorse XFe24 Analyzer (Agilent #1002238-100) using Seahorse Wave Desktop v2.6.0.

### Biofilm quantification

A clear, flat-bottom 96-well plate (Greiner, #M2936) was prepared using either LB or M9 minimal medium supplemented with 0.5% (w/v) glucose, with or without 20 mM itaconate. Each well was inoculated with 1 × 10⁸ CFU of *P. aeruginosa* strains (WT *rpoN*, C218/275A mutant, or Δ*rpoN*) and incubated statically at 37 °C for 24 h (or 48 h for M9 + glucose conditions). Bacterial growth was monitored by measuring optical density at 600 nm (OD₆₀₀) using SkanIt Software 7.0 RE.For biofilm staining, the supernatant was carefully removed, wells were washed and air-dried, and biofilms were fixed with 100% methanol. Biofilms were then stained with 1% crystal violet. After removing the stain, plates were washed, dried, and the biofilm was solubilized in 33% acetic acid. Absorbance at 540 nm was measured using a Varioskan Lux plate reader (Thermo Scientific, #3020-82355).

### Growth curves

Growth assays for bacterial growth curves, a U-bottomed, clear 96-well plate (Greiner Bio-One, 650161) was prepared with 198 µL of M9 minimal media supplemented with glucose an/or succinate, itaconate. Each well was inoculated with 2 µL overnight bacterial culture grown in LB with or without 20mM itaconate, standardized to an OD600nm of 4. Absorbance at 600 nm was read every 10 min for 18 h on the SpectraMax M2 plate reader (Molecular Devices), as the plate incubated at 37C with shaking.

### Swarming motility

Swarming assays were conducted following a previously established protocol ^5^,using LB medium supplemented with 0.5% (w/v) agar. Once solidified, the plates were briefly air-dried at room temperature and spot-inoculated with 2 μL of overnight LB cultures. Plates were incubated face-up in stacks of no more than two at 37 °C for 18 hours. Swarming motility was assessed by measuring the diameter of the swarming zone.

### Protein structure

The three-dimensional structure of the *Pseudomonas aeruginosa* RpoN protein was predicted using AlphaFold 3. PyMOL™ version 3.0.4 (Schrödinger, LLC) software was used to visualize and perform the structural mapping of domains, conserved cysteine residues and mutations.

### Statistical analysis

Experiments in this study were not conducted in a blinded manner. All statistical analyses and graphing were performed using GraphPad Prism 9. Graphical data are presented as mean ± SEM, under the assumption of a normal distribution. For comparisons involving more than two groups, one-way ANOVA followed by post hoc multiple comparisons was used. When analyzing two or more groups over time, two-way ANOVA with post hoc testing was applied. Differences between two groups were assessed using parametric tests (Student’s *t*-test or ANOVA) when data were normally distributed, or nonparametric tests (Mann–Whitney or Kruskal–Wallis) when normality could not be assumed. A two-tailed P value of <0.05 was considered statistically significant. Specific P values, along with the number of independent experiments and replicates, are provided in the figure legends.

## Data Availibility

The MS proteomics data have been deposited at the ProteomeXchange Consortium (http://proteomecentral.proteomexchange.org) via the iProX partner repository (identifier: PXD050510, Subproject ID: IPX0008134002). RNA seq and 13C glucose labeling data as well as other data used in the paper are included in the Supplementary information. Additional raw data are available from the corresponding author upon request.

## Code availability

This work does not include any new code.

### Acknowledgements

This study was funded by NIH grants R01HL170129 (AP), R35HL135800 (AP), T32-5T32DK0076 (GG), S10OD020056 to the Columbia Center for Translational Immunology (CCTI) Flow Cytometry Core.

We thank Luke P Allsopp for providing WT PAO1 and ΔrpoN PAO1 strains. Metabolomics data were acquired at the Calgary Metabolomics Research Facility (CMRF), University of Calgary, and this facility is supported by the International Microbiome Centre and the Canada Foundation for Innovation under Grant CFI-JELF 34986. We thank Dr. Guoan Zhang, Director of the Proteomics and Metabolomics Core Facility at Weill Cornell Medicine for his help with bacterial metabolomics assays.

## Authors Contributions

Conceptualization: A.P, A.Z.B. Methodology: A.Z.B., Y.T.C., T.W.F.L., S.R., A.P. Investigation: A.Z.B.,

Y.T.C., Z.L., G.G., J.M., T.W.F.L., A.T., L.F., L.D, I.L.,C.W. Visualization: A.Z.B., Y.T.C. Funding Acquisition:

A.P. Project Administration: A.P. Supervision: A.P. Writing—original draft: A.P. Writing—review editing: A.P., S.R.and A.Z.B.

## Corresponding author

Correspondence to Dr Alice Prince.

## Competing interests

The authors declare no competing interests.

## Extended figures and legends

**Extended data Fig. 1.**
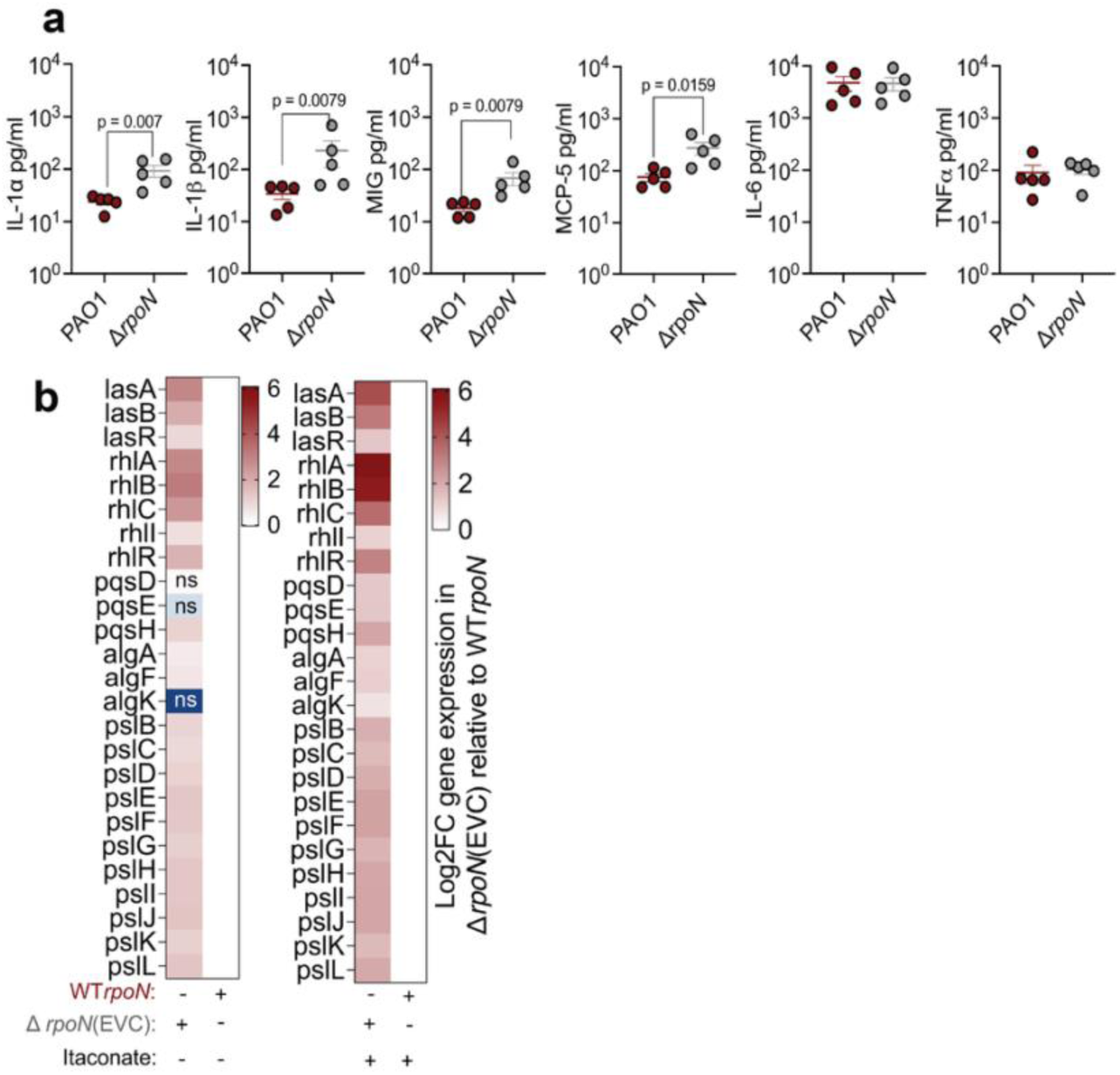
Impact of *rpoN* expression on host and bacterial factors involved in infection. (a) Cytokines measured in BALF at 16h after acute pulmonary infection in C57 BL/NJ6 mice following intranasal inoculation by WT or *ΔrpoN* PAO1 strains. (b) Heat map illustrating the significant effects of itaconate in altering expression of quorum sensing and biofilm associated genes in Δ*rpoN* (EVC) relative to Δ*rpoN* expressing WT*rpoN* grown in LB +/- itaconate, as determined by bulk RNA seq. Data are presented as mean ± s.e.m.; *n* = 2 (a), *n* = 2(b) replicates. Significance is determined by unpaired Mann Whitney U *t*-test (a) and Wald *t* test(b). ns(non-significant).

**Extended data Fig. 2.**
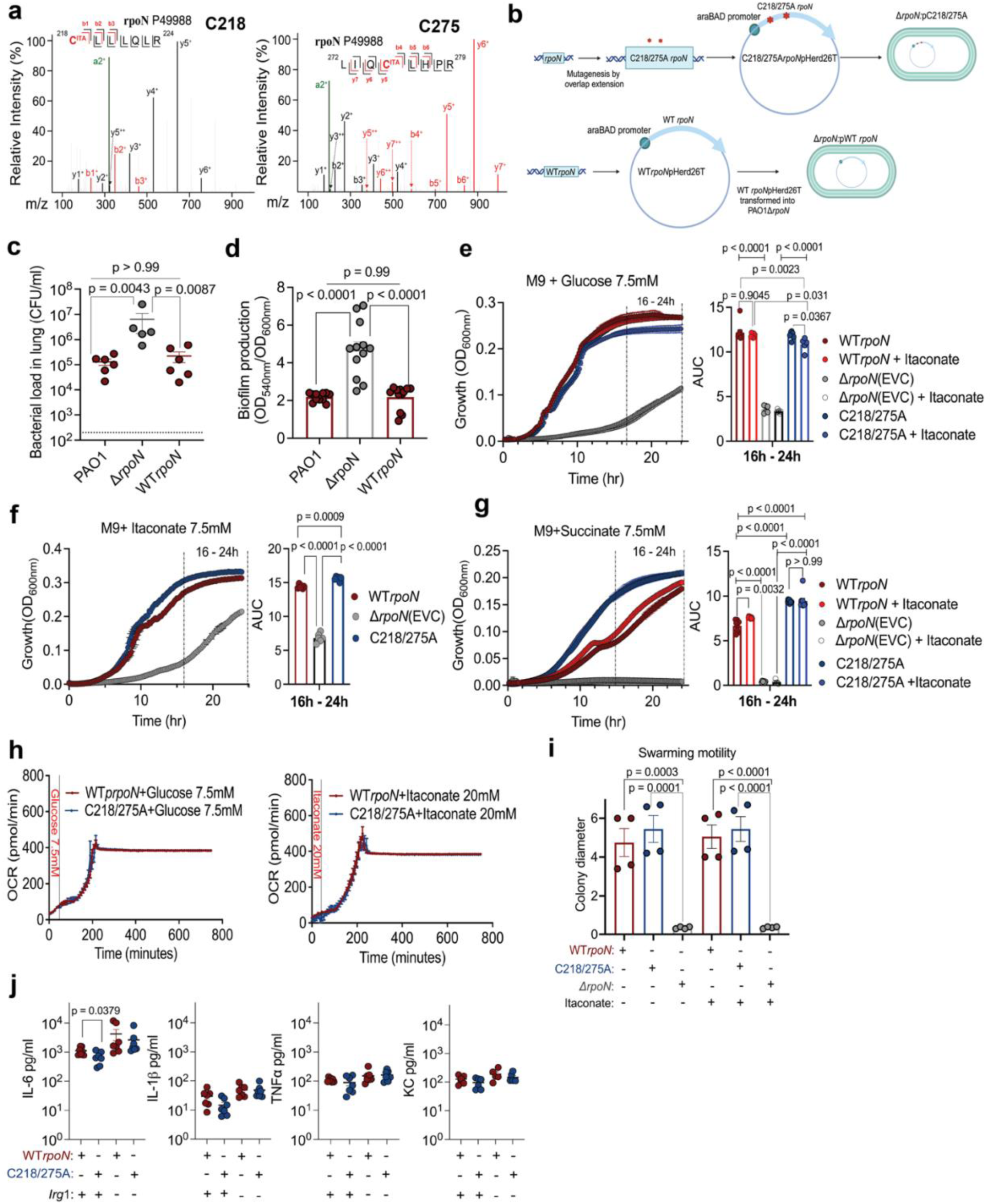
Construction and characterization of strains in which the RpoN cysteines (218/275) are replaced with alanines. (a) MS/MS spectra showing itaconation of cysteines residues 218 and 275 in RpoN. (b) Schematic diagram illustrating generation of plasmid-complementation of *ΔrpoN* PAO1 to generate Δr*poN*:C218/275A mutant and Δ*rpoN*:WT*rpoN* strains. c,d, Complemented Δ*rpoN* expressing WT*rpoN* restores WT PAO1 function. (c) Lung bacterial load post 16 h pulmonary infection. (d) *In vitro* biofilm production (normalized to growth) in LB media at 24 h. e-k, Impact of C to A substitution on RpoN function. e-g, Growth curves of WT*rpoN* or C218/275A or Δ*rpoN* in M9 minimal media supplemented with (e) glucose, (f) itaconate, (g) succinate (e,g -/+ itaconate stress) (statistical significance of differences in growth curve was assessed between 16 and 24 h). (h) Oxygen consumption rate in glucose or itaconate. (i) Swarming motility as measurement of colony diameter in LB agar -/+ itaconate. (j) Cytokines measured in BALF at 16h after acute pulmonary infection in C57 BL/NJ6 WT or Irg1 ^-^/^-^ mice following intranasal inoculation of WT*rpoN* or C218/275A PAO1 strains. Data are presented as mean ± s.e.m.; *n* = 2 (c), *n* = 2 (d), *n* = 2 (e-g), n = 2 (h), n = 2 (i) replicates. Significance is determined by unpaired Mann Whitney U *t*-test (c,j) and One way ANOVA using Tukey’s multiple comparison test with specific two-tailed unpaired *t*-test (d,e,f,g,i).

**Extended data Fig. 3.**
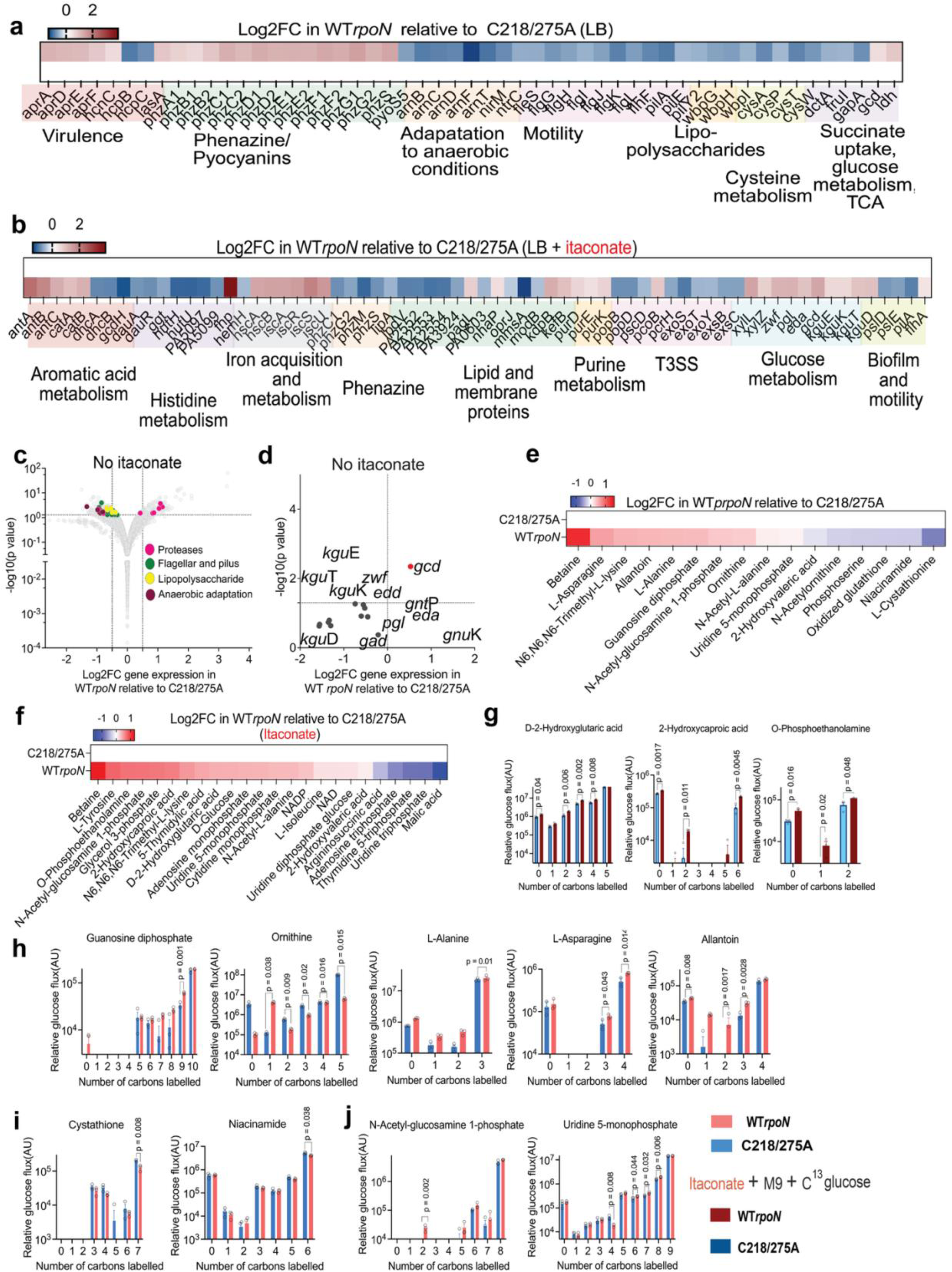
Transcriptomic and metabolic consequences of cysteine to alanine mutations in RpoN. a,d, Bulk RNA seq to determine genes differentially expressed between WT *rpoN* and C218/275A grown in LB media with or without itaconate. Heatmap illustrating significantly altered expression of genes associated with key pathways in C218/275A relative to WT*rpoN* when grown in (a) LB (b) LB+ itaconate. (c) Volcano plot of differentially expressed genes due to C-A substitution. Grey dots represent all genes; colored dots represent genes of specific pathways. Y axis: logarithmic scale, cut off set on Y axis: -log10 (*p* value) ≥ 1.3; cut off set on X axis Log2FC(Fold change) > 1. (d) Volcano plot of differentially expressed genes specific to gluconate or ED pathway due to C-A substitution. Significantly altered total glucose flux (^12^C and ^13^C) due to (e) C to A substitution and (f) itaconate. (g)Increased ^13^C incorporation WT*rpoN* due to itaconate into lipid isotopologues. Cysteine substitution significantly alters ^13^C incorporation into isotopologues associated with various pathways; (h) decreased flux: L-Asparagine, ornithine, betaine, guanosine diphosphate (GDP), (i) increased flux: cystathionine, niacinamide, (j) decreased flux in both groups: UMP, GlcNAc-1-P. Data is presented as mean, *n* = 2 (a-d), *n = 3* (e-j) replicates. Significance is determined by Wald *t* test(a-d) and Unpaired two-tailed Student’s *t*-test (e-j).

**Supplementary Table 3.**
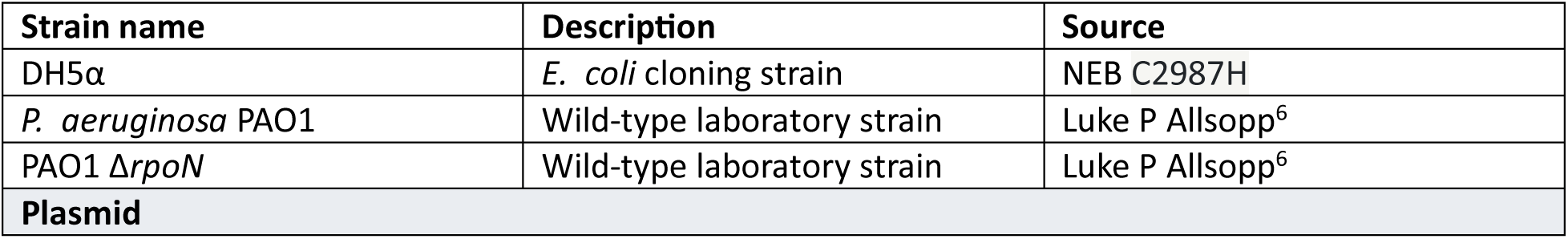
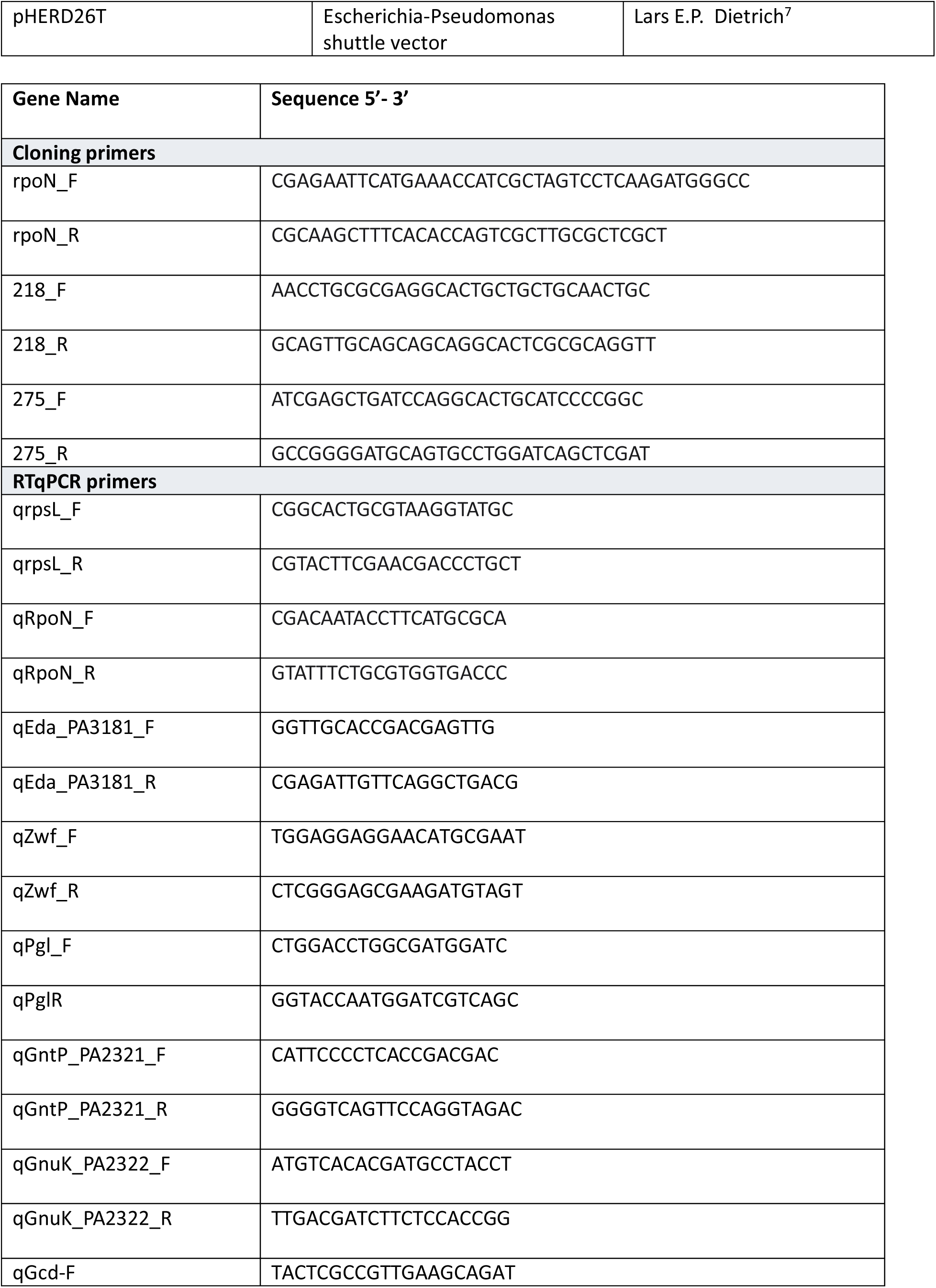
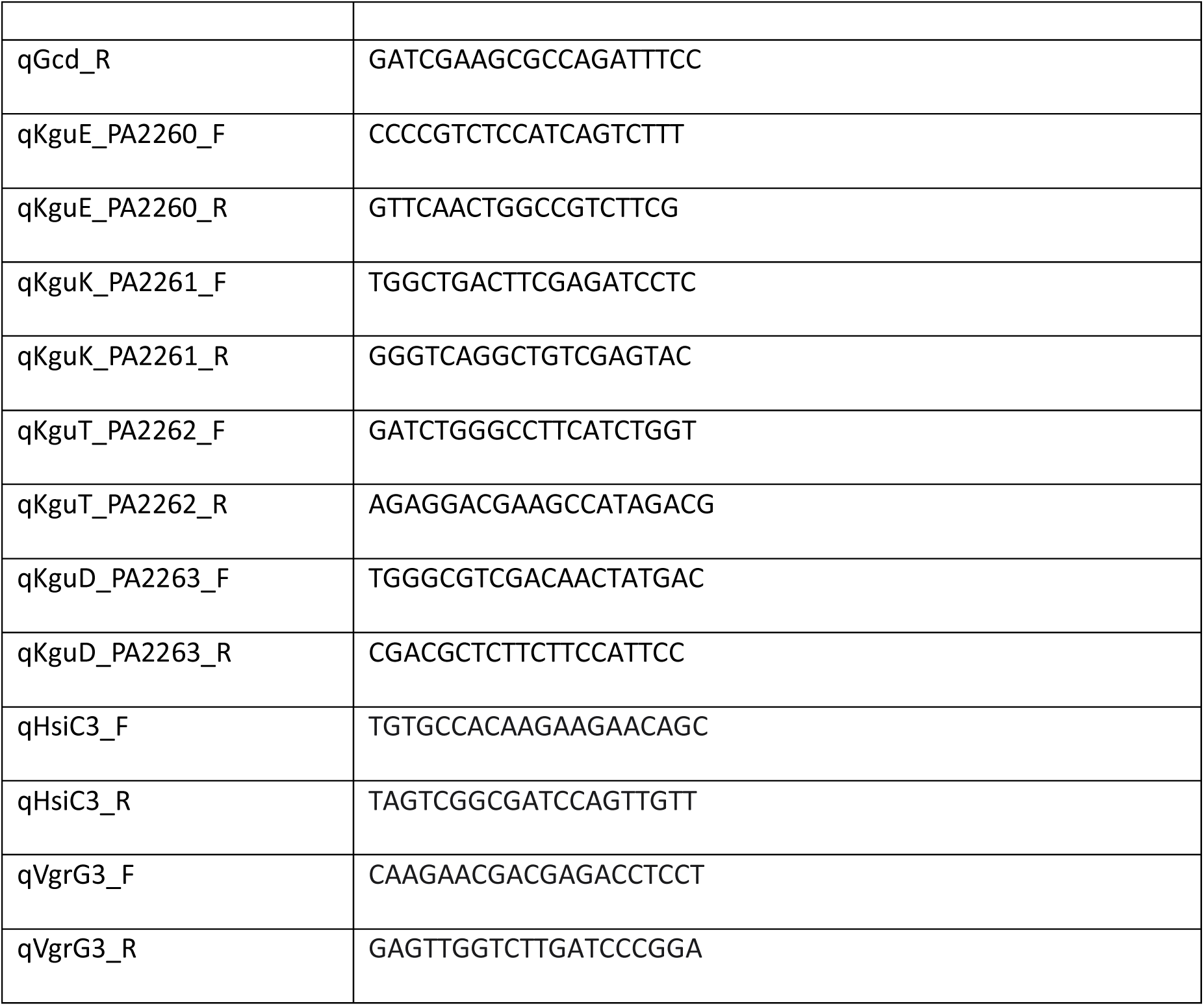
Strains, plasmids and primers used in this study.

## Notes

### Competing Interest Statement

The authors have declared no competing interest.

